# GWAS Meta-Analysis of Neuroticism (N=449,484) Identifies Novel Genetic Loci and Pathways

**DOI:** 10.1101/184820

**Authors:** Mats Nagel, Philip R Jansen, Sven Stringer, Kyoko Watanabe, Christiaan A de Leeuw, Julien Bryois, Jeanne E Savage, Anke R Hammerschlag, Nathan Skene, Ana B Muñoz-Manchado, the 23andMe Research Team, Sten Linnarsson, Jens Hjerling-Leffler, Tonya JH White, Henning Tiemeier, Tinca JC Polderman, Patrick F Sullivan, Sophie van der Sluis, Danielle Posthuma

## Abstract

Neuroticism is an important risk factor for psychiatric traits including depression^1^, anxiety^2,3^, and schizophrenia^4–6^. Previous genome-wide association studies^7–12^ (GWAS) reported 16 genomic loci^10–12^. Here we report the largest neuroticism GWAS meta-analysis to date (N=449,484), and identify 136 independent genome-wide significant loci (124 novel), implicating 599 genes. Extensive functional follow-up analyses show enrichment in several brain regions and involvement of specific cell-types, including dopaminergic neuroblasts (*P*=3×10^-8^), medium spiny neurons (*P*=4×10^-8^) and serotonergic neurons (*P*=1×10^-7^). Gene-set analyses implicate three specific pathways: neurogenesis (*P*=4.4×10^-9^), behavioural response to cocaine processes (*P*=1.84×10^-7^), and axon part (P=5.26×10^-8^). We show that neuroticism’s genetic signal partly originates in two genetically distinguishable subclusters^13^ (*depressed affect* and *worry*, the former being genetically strongly related to depression, *rg*=0.84), suggesting distinct causal mechanisms for subtypes of individuals. These results vastly enhance our neurobiological understanding of neuroticism, and provide specific leads for functional follow-up experiments.

---

The neuroticism meta-analysis comprised data from the UK Biobank Study (UKB, full release^14^; N=372,903; **Online Methods; Supplementary Fig. 1**), 23andMe, Inc.^15^ (N=59,206), and the Genetics of Personality Consortium (GPC1^9^; N=17,375; **Online Methods**, N**=**449,484 in total). In all samples, neuroticism was measured through (digital) questionnaires (**Online Methods; Supplementary Information**). SNP associations were meta-analyzed using METAL^16^, weighted by sample size (**Online Methods)**. The quantile-quantile (Q-Q) plot of the genome-wide meta-analysis on 449,484 subjects and 14,978,477 SNPs showed high inflation (λ=1.65) and mean *χ*^2^ statistic (1.91) **(Fig. 1a**; **Supplementary Table 1)**. The LD score regression (LDSC)^17,18^ intercept (1.02; SE=0.01) was consistent with inflation due to true polygenicity and large sample size. The LDSC SNP-based heritability (*h^2^ SNP*) of neuroticism was 0.100 (SE=0.003).

**Fig. 1.**
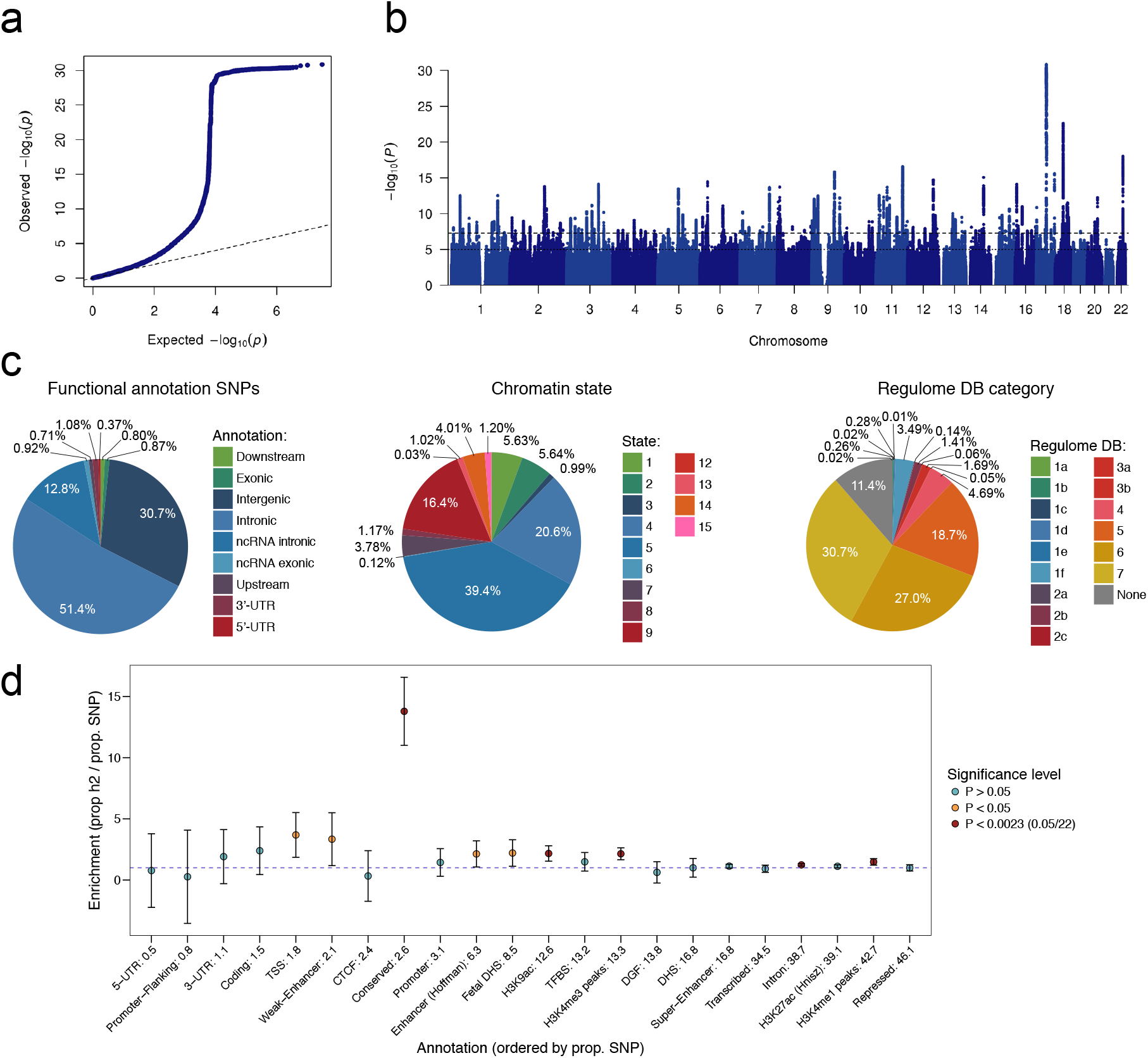
SNP-based associations with neuroticism in the GWAS meta-analysis. (a) Quantile-quantile plot of the SNP-based associations with neuroticism. (b) Manhattan plot showing the -log10 transformed *P*-value of each SNP on the y-axis and base pair positions along the chromosomes on the x-axis. The dashed line indicates genome-wide significance (*P*<5×10^-8^), the dotted line the threshold for suggestive associations (*P*<1×10^-5^). (c) Pie charts showing the distribution of functional consequences of SNPs in linkage disequilibrium (LD) with genome-wide significant lead SNPs in the meta-analysis, the minimum chromatin state across 127 tissue and cell types and the distribution of regulome DB score, a categorical score between 1a and 7, indicating biological evidence of a SNP being a regulatory element, with a low score denoting a higher likelihood of being regulatory. (d) Heritability enrichment of 22 functional SNP annotations calculated with stratified LD score regression.

**Table 1.**
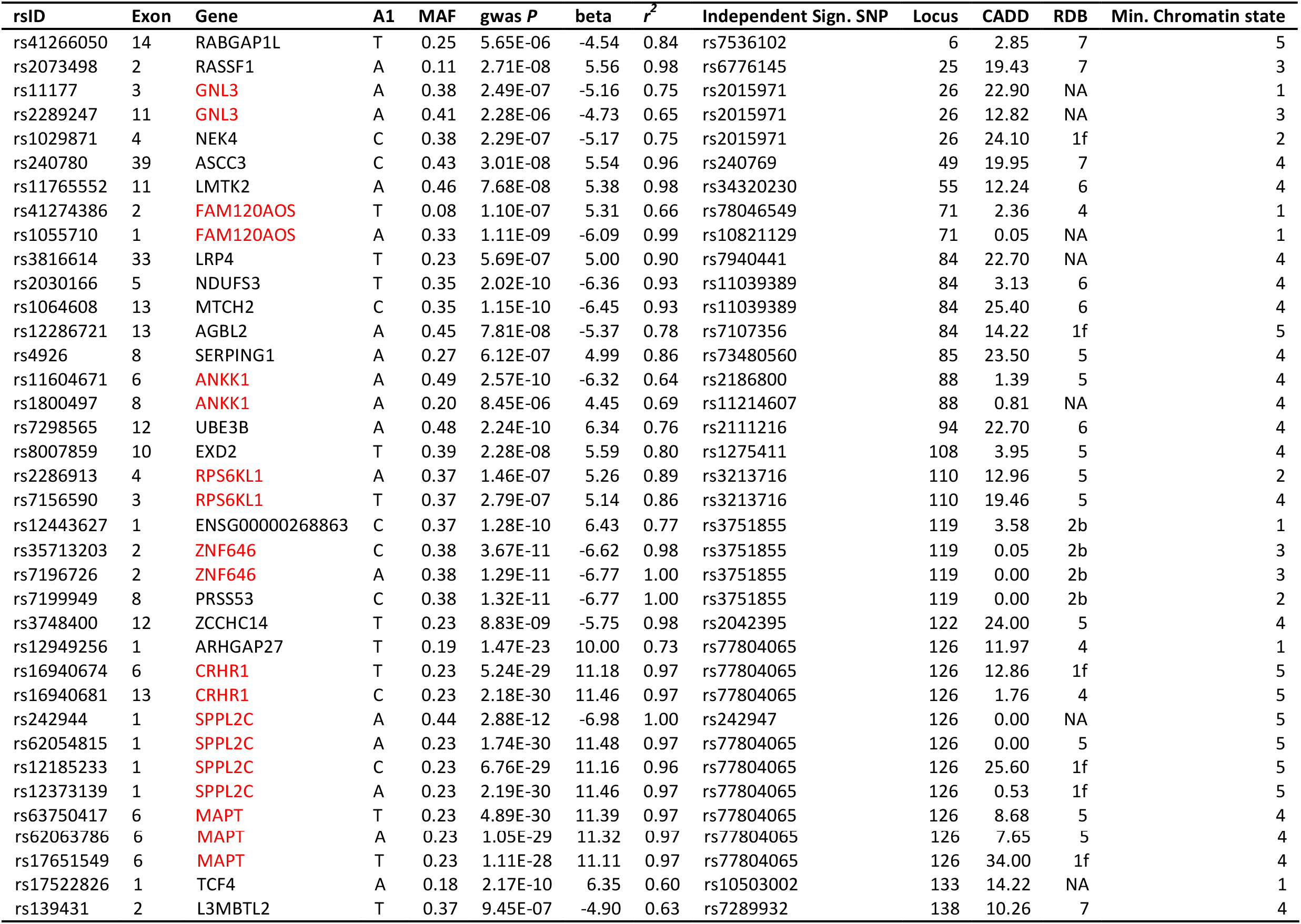
Exonic non-synonymous (ExNS) variants in the genomic loci associated with neuroticism and in LD (*r^2^*>0.6) with one of the independent GWS SNPs. CADD: CADD score; rdb: regulome DB score; MAF: minor allele frequency; Z-score: z-score from the GWAS meta-analysis in METAL. Results are reported on hg19 coordinates (NCBI b37). Genes containing multiple ExNS are annotated in red.

The GWAS meta-analysis identified 9,745 genome-wide significant (GWS) SNPs (*P*<5×10^-8^), of which 157 and 2,414 were located in known associated inversions on chromosomes 8 and 17^10–12^, respectively (**Supplementary Table 2**; **Fig. 1b**; **Supplementary Fig. 2**). FUMA^19^, a tool to functionally map and annotate GWAS results (**Online Methods**), extracted 170 independent lead SNPs (158 novel; see **Supplementary information** for definition of lead SNPs), which mapped to 136 independent genomic loci (124 novel; **Online Methods**; **Supplementary information; Supplementary Table 3-8**). Of all lead SNPs, 4 were in exonic, 88 in intronic, and 52 in intergenic regions. Of the 17,794 SNPs in high LD with one of the independent significant SNPs (see **Supplementary information** for definition of independent significant SNPs), most were intronic (9,147: 51,4%) or intergenic (5,460: 30,7%), and 3.8% was annotated as potentially having a functional impact, with 0.9% (155 SNPs) being exonic (**Fig. 1c**, **Supplementary Table 9**; see **Supplementary Tables 10-11** for an overview of chromatin state and regulatory functions of these SNPs). Of these, 37 were exonic nonsynonymous (ExNS) (**Table 1, Supplementary Table 12**). The highest CADD score (34) of ExNS SNPs was for rs17651549, in exon 6 of *MAPT*, with a GWAS *P*-value of 1.11´10^-28^, in high LD with the lead SNP in that region (*r*^*2*^=0.97). rs17651549 is a missense mutation leading to an Arginine to Tryptophan change with allele frequencies matching the inversion in that region. The ancestral allele C is associated with a lower neuroticism score (see **Table 1** and **Supplementary Table 12** for a detailed overview of all functional variants in genomic risk loci).

Stratified LDSC^20^ (**Online Methods**), showed significant enrichment for *h*^2^ of SNPs located in conserved regions (enrichment=13.79, *P*=5.14×10^-16^), intronic regions (enrichment=1.27, *P*=1.27×10^-6^), and in H3K4me3 (enrichment=2.14, *P*=1.02×10^-5^) and H3K9ac regions (enrichment=2.17, *P* = 3.06×10^-4^) (**Fig. 1d**; **Supplementary Table 13).**

Polygenic scores (PGS) calculated using PRSice^21^ (clumping followed by *P*-value thresholding) and LDpred^22^ in three randomly drawn hold-out samples (UKB only, N=3,000 each; **Online Methods**), explained up to 4.2% (*P*=1.49×10^-30^) of the variance in neuroticism **(Supplementary Fig. 3; Supplementary Table 14**).

**Fig 3.**
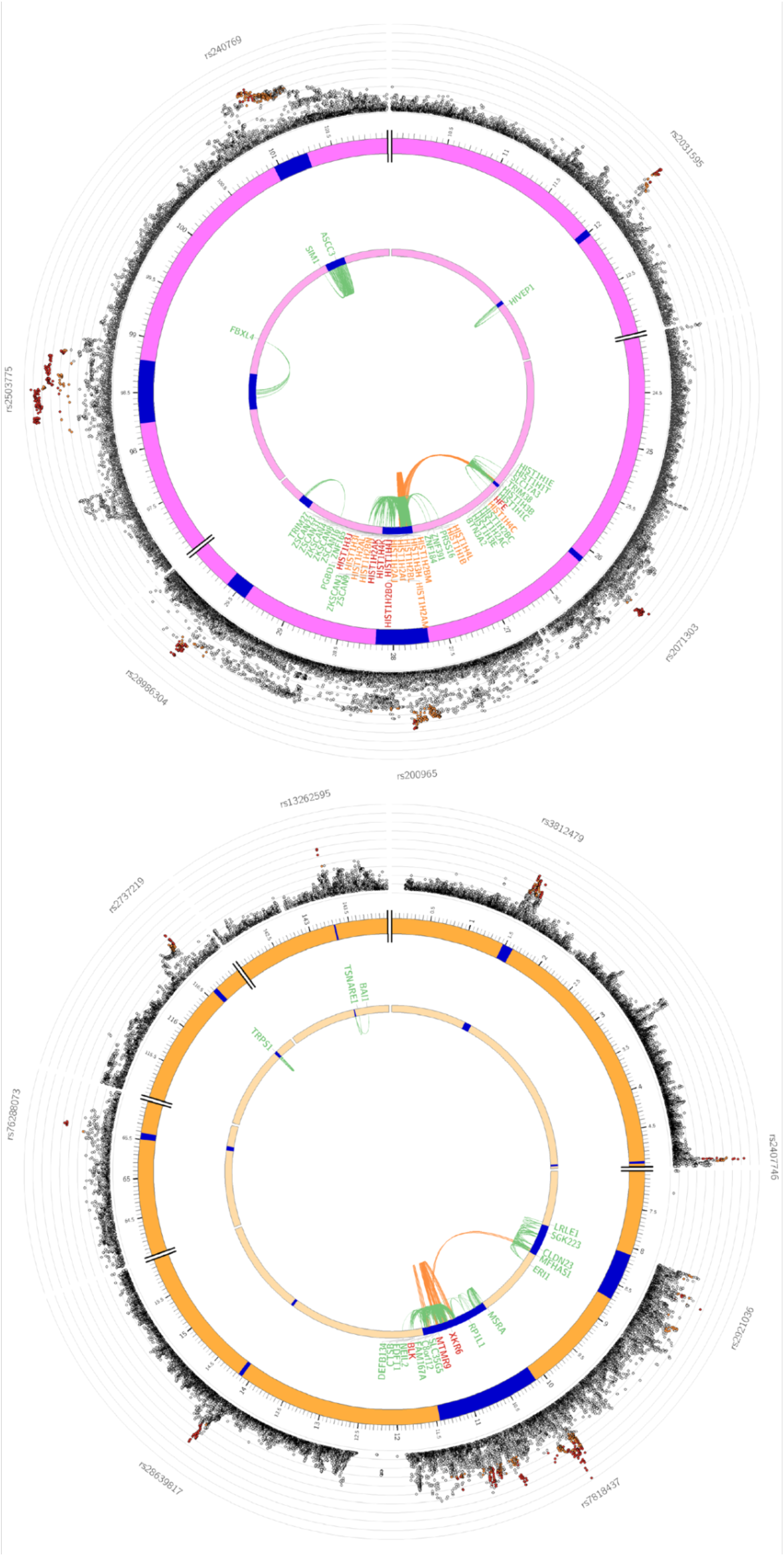
Genomic risk loci, eQTL associations and chromatin interaction for chromosome 6 and 8, containing cross-locus interactions. Circos plot showing genes on (a) chromosome 6 and (b) chromosome 8 that were implicated by the genomic risk (blue areas) loci by chromatin interaction (CTI; orange), eQTL (green) or implicated by both eQTL and CTI mapping (red). The outer layer shows a Manhattan plot containing the -log10 transformed *P*-value of each SNP in the GWAS meta-analysis. Empty regions in the Manhattan plot layer indicate regions where no SNPs with *P*<0.05 are situated.

We used four strategies to link our SNP results to genes: positional, eQTL, and chromatin interaction mapping (**Online Methods**) and genome-wide gene-association analysis (GWGAS; MAGMA^23^). GWGAS evaluates the joint association effect of all SNPs within a gene yielding a gene-based *P*-value. Based on our meta-analytic results, 283 genes were implicated through positional mapping, 369 through eQTL-mapping, and 119 through chromatin interaction-mapping (**Fig. 2a; Supplementary Table 15**). GWGAS identified 336 GWS genes (*P*<2.75×10^-6^, **Figs. 2b-c**; **Supplementary Table 16, Supplementary information**), of which 203 overlapped with genes implicated by FUMA, resulting in 599 unique neuroticism-related genes. Of these, 50 were implicated by all four methods, of which 49 had chromatin interaction and eQTL associations in the same tissue/cell type (**Fig. 2a, Supplementary Table 15**).

**Fig. 2.**
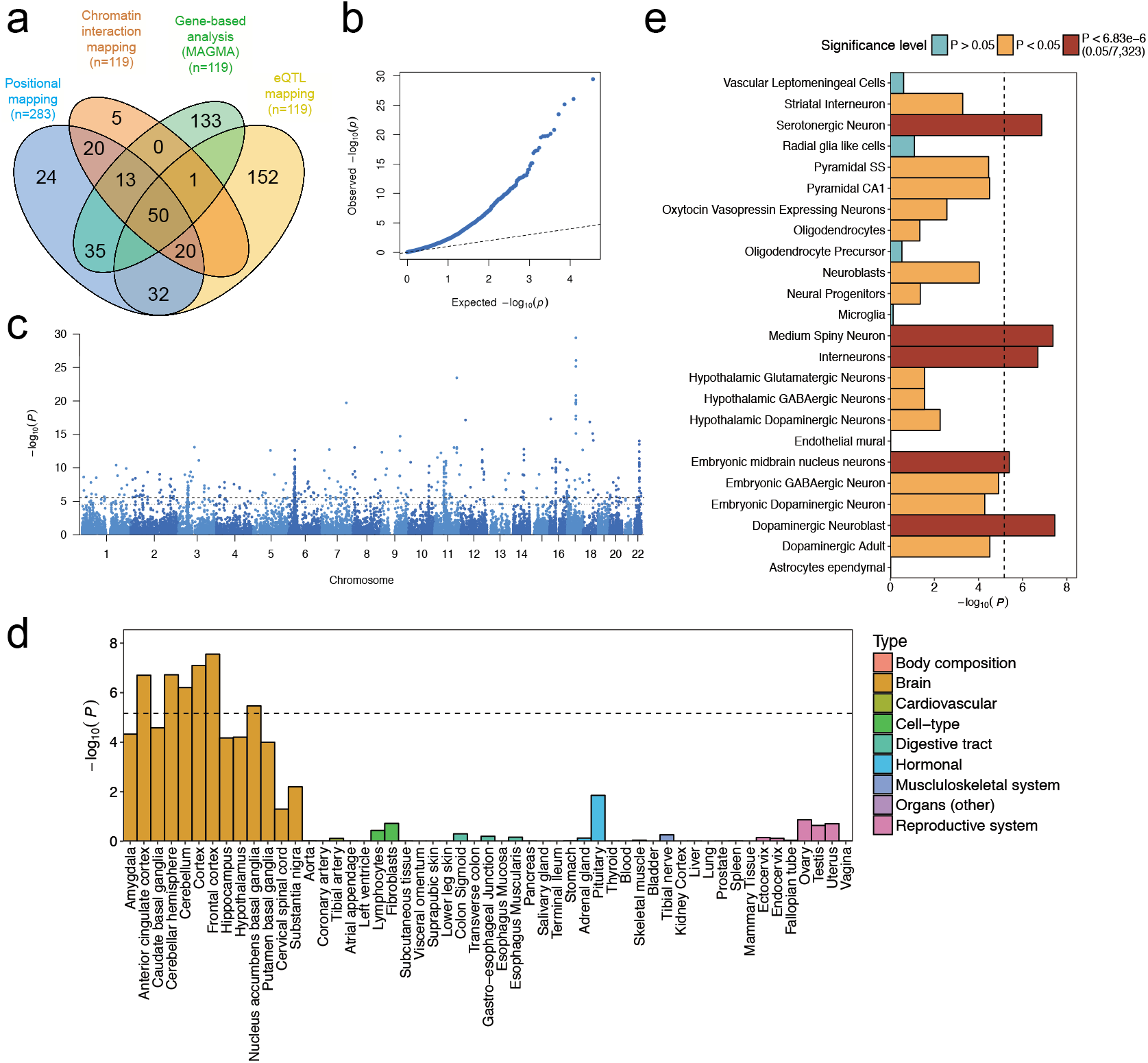
Mapping of genes and tissue- and cell expression profiles. a) Venn diagram showing overlap of genes implicated by positional mapping, eQTL mapping, chromatin interaction mapping, and gene-based genome-wide association (GWGAS). (b) Quantile-quantile plot of the GWGAS. (c) Manhattan plot of the genome-wide gene-based association analysis (GWGAS) on neuroticism. The y-axis shows the -log10 transformed *P*-value of each gene, and the chromosomal position (start position) on the x-axis. The dashed line indicates the threshold for genome-wide significance of the gene-based test (*P*< 2.76×10^-6^; 0.05/18,128), and the dotted line indicates the suggestive threshold (*P*<2.76×10^-5^; 0.5/18,128). (d) Gene expression profiles of identified genes for 53 tissue types. Expression data were extracted from the Genotype-Tissue Expression (GTEx) database. Expression values (RPKM) were log2 transformed with pseudocount 1 after winsorization at 50 and averaged per tissue. Gene-set tests for tissue expressions were calculated using MAGMA (Online Methods). (e) Enrichment of genetic signal for neuroticism in 24 brain cell types. The dashed line indicates the Bonferroni-corrected significance threshold (*P*=0.05/7,323=6.83´10^-6^).

19 of the 119 genes implicated through chromatin interaction mapping are especially interesting as they are implicated via interactions between two independent GWS genomic risk loci. There are several chromatin interactions in 7 tissue types (aorta, hippocampus, left ventricle, right ventricle, liver, spleen, pancreas) across two risk loci on chromosome 6 (**Fig. 3a**). Two genes are located in locus 45 and are mapped by chromatin interactions from risk locus 46 (*HFE* and *HIST1H4C*), and another 16 genes are coding histones in locus 46 and are mapped by interactions from locus 45 (**Supplementary Table 15**). *XKR6* is located on chromosome 8 in risk locus 61, and is implicated by chromatin interactions in 5 tissue types (aorta, left ventricle, liver, pancreas and spleen) including cross locus interactions from locus 60 (**Fig. 3b**; **Supplementary Table 15**). This gene is also mapped by eQTLs in blood and transformed fibroblasts. Out of the 19 genes mapped by two loci, 4 are located outside of the risk loci (*HIST1H2AI, HIST1H3H, HIST1H2AK* and *HIST1H4L*), and 7 are also implicated by eQTLs in several tissue types (*HFE* in adipose subcutaneous, aorta, esophagus muscularis, lung, tibial nerve, sub-exposed skin and thyroid; *HIST1H4J* in blood and adrenal gland; and *HIST1H4K, HIST1H2AK, HIST1H2BO* and *XKR6* in blood).

Gene-based *P*-values were used for gene-set analysis in MAGMA^23,26^, testing 7,246 predefined gene-sets derived from MsigDB^24^, gene expression profiles in 53 tissue types obtained from the GTEx Project^25^, and 24 cell-type specific expression profiles using RNAseq information^26^ **(Online Methods)**. Neuroticism was significantly associated with genes predominantly expressed in 11 brain tissue types (**Fig. 2d**; **Supplementary Table 17-18)** and with 7 gene ontology (GO) gene-sets, with the strongest association for neurogenesis (*P*=0.0004) and neuron differentiation (*P*=0.002) (**Supplementary Table 17)**. Conditional gene-set analyses (**Online Methods**) suggested that 3 of the 7 gene-sets (neurogenesis, *P*=4.4×10^-9^; behavioral response to cocaine, *P*=1.84×10^-7^; axon part, P=5.26×10^-8^) had largely independent associations, implying a role in neuroticism (**Supplementary Table 19**). Conditional analyses of the tissue-specific expression ascertained general involvement of (frontal) cortex expressed genes (**Supplementary Table 20**; **Supplementary Fig. 4**).

Cell type specific gene-set analysis showed significant association with genes expressed in multiple brain cell types **(Fig. 2e**; **Supplementary Table 21**), with dopaminergic neuroblasts (*P*=3×10^-8^), medium spiny neurons (*P*=4×10^-8^) and serotonergic neurons (*P*=1×10^-7^) showing the strongest associations, and conditional analysis indicated that these three cell types were also independently associated with neuroticism.

Aiming to further specify neuroticism’s neurobiological interpretation, we compared the genetic signal of the full neuroticism trait to that of two genetically distinguishable neuroticism subclusters *depressed affect* and *worry*^13^ (**Online Methods**). As a validation of the *depressed affect* dimension, we also compare with GWAS results for depression. GWA analyses of the subclusters were conducted on the UKB-data only (dictated by item-level data availability; **Online Methods**; *depressed affect*, N=357,957; *worry*, N=348,219). For depression, our metaanalysis comprised data from the UKB^14^ (N=362,696; **Supplementary Fig. 5**), 23andMe^15^ (N=307,354), and the Psychiatric Genetics Consortium (PGC^27^; N=18,759) (total N=688,809, not previously published; *r_g_* between samples: 0.61-0.80; **Online Methods**; **Supplementary Table 22, Supplementary Information**). Genetic correlations of neuroticism with all three phenotypes were considerable (depression: *r*g=0.79; *depressed affect*: *r*g=0.88, *worry*: *r*g=0.87; **Supplementary Table 23**).

The subclusters showed notable differences in genetic signal (e.g., exclusive GWS associations on chromosomes 2 and 19 for *depressed affect*, and chromosomes 3 and 22 for *worry*; **Supplementary Figs. 6-12**; **Supplementary Tables 24-26**). Of the 136 genetic loci associated with neuroticism, 32 were also GWS for *depressed affect* (7 shared with depression) but not for *worry*, and 26 were also GWS for *worry* (3 shared with depression) but not for *depressed affect* (**Supplementary Table 27**; **Supplementary Fig. 12**). These results were mirrored by gene-based analyses (**Supplementary information**; **Supplementary Tables 28-30**; **Supplementary Fig. 13**), suggesting that part of neuroticism’s genetic signal originates specifically in one of the two subclusters, possibly implicating different causal genetic mechanisms.

To test specificity of the gene-sets implicated in neuroticism in the conditional analyses, we repeated the analyses, but now corrected for *depressed affect*, and *worry* scores, respectively (**Supplementary Table 31**; **Supplementary Fig. 14**). The association with ‘axon-part’ was markedly lower after correction for *worry* scores (uncorrected *P*=5.26×10^-8^; corrected for *depressed affect P*=2.42×10^-6^; corrected for *worry P*=.0013), suggesting that the involvement of ‘axon-part’ in neuroticism originates predominantly from the *worry*-component.

To examine the genetic correlational pattern of neuroticism, and to compare it to the patterns observed for depression, *depressed affect* and *worry*, we used LDSC to calculate genetic correlations with 35 traits for which large-scale GWAS summary statistics were available (**Supplementary Table 32**; **Online Methods**). We observed 11 Bonferroni-corrected significant genetic correlations for neuroticism (α=0.05/(4´35); *P*<3.6×10^-4^) (**Fig. 4; Supplementary Table 33)**, covering previously reported psychiatric traits (*rg* range: .20-.82) and subjective well-being (*rg*= -.68). These correlations were supported by enrichment of neuroticism genes in sets of genes previously implicated in psychiatric traits (**Supplementary Table34**). The *rg*’s of depression and *depressed affect* strongly mirrored eachother (correlation between their *rg*’s is *r*=.98; **Supplementary information**), validating the *depressed affect* cluster. The correlational patterns for *depressed affect* and *worry* were markedly different and sometimes antipodal, with the genetic signal of the full neuroticism trait being a blend of both.

**Fig. 4.**
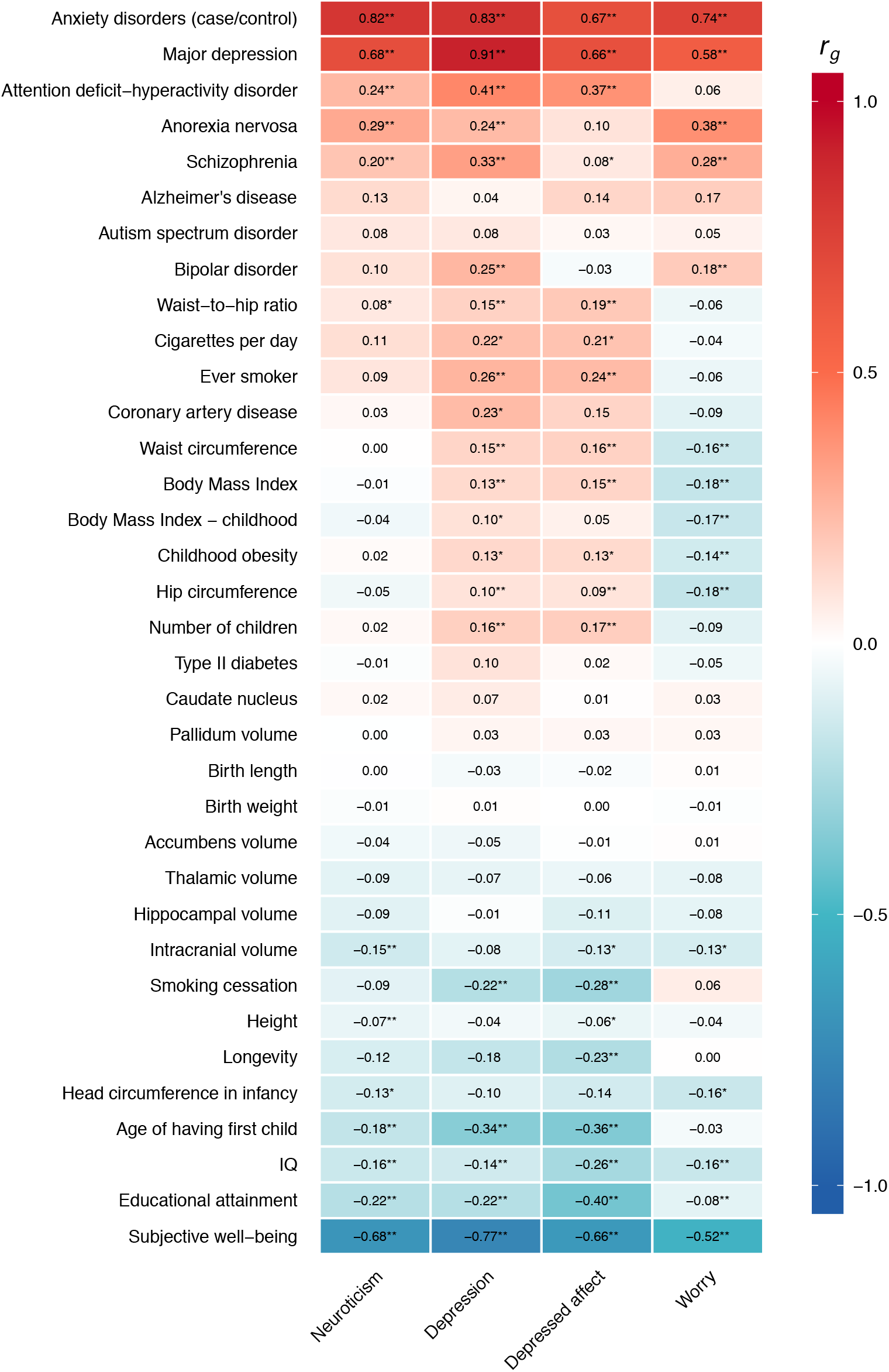
Genetic correlations between neuroticism and other traits. Genetic correlations of neuroticism, depression, *depressed* affect and worry with various traits and diseases. LD score regression (Online Methods) tested genome-wide SNP associations for the neuroticism score against previously published results for 35 neuropsychiatric outcomes, antropometric and health-related traits, and brain morphology (Supplementary Table 32-33).

In conclusion, we identified 119 novel genetic loci for neuroticism. Extensive functional annotations highlighted several genes being implicated through multiple routes. We demonstrated the involvement of specific neuronal cell types and three independently associated genetic pathways, and established the genetic multidimensionality of the neuroticism phenotype, and its link with depression. The current study provides new leads, and testable functional hypotheses for unraveling the neurobiology of neuroticism, its subtypes, and genetically associated traits.

## URLs

<u>http://ukbiobank.ac.uk</u>

<u>http://ctg.cncr.nl/software/magma</u>

<u>http://software.broadinstitute.org/gsea/msigdb/collections.jsp</u>

<u>http://genome.sph.umich.edu/wiki/METAL_Program</u>

<u>https://github.com/bulik/ldsc</u>

<u>http://fuma.ctglab.nl</u>

## Acknowledgements

This work was funded by The Netherlands Organization for Scientific Research (NWO Brain & Cognition 433-09-228, NWO MagW VIDI 452-12-014, NWO VICI 435-14-005 and 453-07-001, 645-000-003), Sophia Foundation for Scientific Research (SSWO, grant nr: S-1427). JB was funded by the Swiss National Science Foundation. Analyses were carried out on the Genetic Cluster Computer, which is financed by the Netherlands Scientific Organization (NWO: 480-05-003), by the VU University, Amsterdam, the Netherlands, and by the Dutch Brain Foundation, and is hosted by the Dutch National Computing and Networking Services SurfSARA. This research has been conducted using the UK Biobank Resource (application number 16406). We would like to thank the participants, including the 23andMe customers who consented to participate in research, and researchers who collected and contributed to the data.

## Author Contributions

S.vd.S and D.P. conceived the study. M.N. and P.R.J. performed the analyses. S. St. performed the quality control on the UK Biobank data and wrote a pipeline to facilitate data processing. K.W. constructed the tool for biological annotation and ran the analyses. J.B. and P.F.S. performed the single-cell gene-expression analysis. M.N., P.R.J., S.vd.S and D.P. wrote the paper. All authors discussed the results and commented on the paper.

## 23andMe Research Team contributors

Michelle Agee, Babak Alipanahi, Adam Auton, Robert K. Bell, Katarzyna Bryc, Sarah L. Elson, Pierre Fontanillas, Nicholas A. Furlotte, David A. Hinds, Bethann S. Hromatka, Karen E. Huber, Aaron Kleinman, Nadia K. Litterman, Matthew H. McIntyre, Joanna L. Mountain, Elizabeth S. Noblin, Carrie A.M. Northover, Steven J. Pitts, J. Fah Sathirapongsasuti, Olga V. Sazonova, Janie F. Shelton, Suyash Shringarpure, Chao Tian, Joyce Y. Tung, Vladimir Vacic, and Catherine H. Wilson.

## Author Information

Reprints and permissions information is available at www.nature.com/reprints. The authors declare no competing financial interest. Correspondence and resquests for materials should be addressed to.

## Online methods

### Samples

#### UK Biobank

The UK Biobank (UKB) Study is a major data resource, containing genetic and a wide range of phenotypic data of ~500,000 participants aged 40-69 at recruitment^14^. We used data released in July 2017, and selection (discussed below) resulted in final sample sizes of N=372,903 and N=362,696 individuals for neuroticism and depression, respectively (**Supplementary information)**. The UKB received ethical approval from the National Research Ethics Service Committee North West–Haydock (reference 11/NW/0382), and all study procedures were performed in accordance with the World Medical Association for medical research. The current study was conducted under UKB application number 16406.

#### 23andMe

23andMe, Inc. is a large personal genomics company that provides genotype and health-related information to customers. For the neuroticism meta-analysis, we used neuroticism GWAS summary statistics from a subset of 23andMe research participants (N=59,206), described in more detail elsewhere^10^. For our depression meta-analysis, we used depression GWAS summary statistics from a subset of 23andMe research participants (N=307,354), described in detail elsewhere^28^. All included participants provided informed consent and were of European ancestry, and related individuals were excluded. Online data collection procedures were approved by the Ethical & Independent Review Services (E&I Review), an AAHRPP-accredited private institutional review board (http://www.eandireview.com).

#### Genetics of Personality Consortium

The Genetics of Personality Consortium (GCP) is a large body of cooperation concerning GWAS on personality. We used summary statistics of neuroticism from the first GCP personality meta-analysis (GPC1, http://www.tweelingenregister.org/GPC/)^9^, on 10 discovery cohorts (SardiNIA, NTR/NESDA, ERF, SAGE, HBCS, NAG, IRPG, QIMR, LBC1936, BLSA, EGPUT), including in total N=17,375 participants of European descent. All included studies were approved by local ethic committees, and informed consent was obtained from all participants.

#### Psychiatric Genetics Consortium

The Psychiatric Genetics Consortium (PGC) unites investigators worldwide to conduct genetic meta- and mega-analyses for psychiatric disorders. We used summary statistics from the latest published PGC meta-analysis on depression (http://www.med.unc.edu/pgc/results-and-downloads)^27^, which included data from 8 cohorts (Bonn/Mannheim, GAIN, GenRED, GSK, MDD2000, MPIP, RADIANT, STAR*D), covering N=18,759 participants of European descent. All included studies were approved by local ethic committees, and informed concent was obtained from all participants.

### Phenotype assessment – Neuroticism

#### UK Biobank

Neuroticism was measured with 12 dichotomous (yes/no) items of the Eysenck Personality Questionnaire Revised Short Form (EPQ-RS^29^, using a touchscreen-questionnaire at the UKB assessment centers (**Supplementary Information 1.1**). Participants with valid responses to <10 items were excluded from analyses. A weighted neuroticism sum score was calculated by adding up individual valid item responses, and dividing that sum by the total number of valid responses. Scores on 4 EPQ-RS items (i.e., “Do you often feel lonely?”, “Do you ever feel ‘just miserable’ for no reason?”, “Does your mood often go up and down?”, and “Do you often feel ‘fed-up’?”) were summed to obtain scores for the cluster *depressed affect*. Similarly, scores on 4 other EPQ-RS items (i.e., “Are you a worried?”, “Do you suffer from nerves?”, “Would you call yourself a nervous person?”, and “Would you call yourself tense or highly strung”) were summed to obtain scores for the cluster *worry*. In the item-cluster analyses, only participants with complete scores on all 4 items were included, resulting in N=357,957 and N=348,219 for *depressed affect* and *worry*, respectively.

#### 23andMe

Neuroticism was operationalized as of the sum of 8 neuroticism items (5-point Likert scale; ‘Disagree strongly’ to ‘Agree strongly’) from the Big Five Inventory (BFI^30,31^), as obtained in an online survey. Only participants with valid responses to all items were included in the analyses (**Supplementary Information 1.2**).

#### Genetic Personality Consortium

All 10 cohorts included in the first meta-analysis of the GPC used sums of the scores on 12 items (5-point Likert scale; ‘Strongly disagree’ to ‘Strongly agree’) of the NEO-FFI^32^ to measure neuroticism. If <4 item scores were missing, data on invalid items were imputed by taking an individual’s average score on valid items. Participants were excluded from analyses if they had invalid scores on >3 items^9^ (**Supplementary Information 1.3**).

### Phenotype assessment – Depression

#### UK Biobank

Depression was operationalized by adding up the scores on two continuous items (“Over the past two weeks, how often have you felt down, depressed or hopeless?”, “Over the past two weeks, how often have you had little interest or pleasure in doing things?”; both evaluated on a 4-point Likert scale; ‘Not at all’ to ‘Nearly every day’), resulting in a continuous depression score (as used previously^12^). Only participants with scores on both items were included in the analyses, resulting in N=362,696 (**Supplementary Information 1.4**).

#### 23andMe

This concerns a case-control sample. Four self-report survey items were used to determine case-control status. Cases were defined as replying affirmatively to at least one of these questions, and not replying negatively to previous ones. Controls replied negatively to at least one of the questions, and did not report being diagnosed with depression on previous ones (**Supplementary Information 1.5**).

#### Psychiatric Genetics Consortium

This concerns a case-control sample. Cases had a DSM-IV lifetime (sometimes (early onset) recurrent) major depressive disorder (MDD) diagnosis, either established through structured diagnostic interviews or clinician-administered DSM-IV checklists. Most cases were ascertained from clinical sources, while controls were randomly selected from population resources and screened for lifetime history of MDD^27^ (**Supplementary Information 1.6**).

### Genotyping and imputation

#### UK Biobank – Neuroticism

We used genotype data released by the UKB in July 2017. The genotype data collection and processing are described in detail by the responsible UKB group^14^. In short, 489,212 individuals were genotyped on two customized SNP arrays (the UK BiLEVE Axiom array (N=50,520) and UK Biobank Axiom array (N=438,692)), covering 812,428 unique genetic markers (95% overlap in SNP content). After quality control procedures^14^, 488,377 individuals and 805,426 genotypes remained. Genotypes were phased and imputed by the coordinating team to approximately 96 million genotypes using a combined refence panel including the Haplotype Reference Consortium and the UK10K haplotype panel. Imputed and quality controlled genotype data was available for 487,422 individuals and 92,693,895 genetic variants. As recommended by the UKB team, variants imputed from the UK10K reference panel were removed from the analyses due to technical errors in the imputation process.

In our analyses, only individuals from European descent (based on genetic principal components) were included. Therefore principal components from the 1000 Genomes reference populations^33^ were projected onto the called genotypes available in UK Biobank. Subjects were identified as European if their projected principal component score was closest (based on the Mahalanobis distance) to the average score of the European 1000 Genomes sample^34^. European subjects with a Mahalanobis distance > 6 S.D. were excluded. In addition, participants were excluded based on withdrawn consent, UKB provided relatedness (subjects with most inferred relatives, 3^rd^ degree or closer, were removed until no related subjects were present), discordant sex, sex aneuploidy. After selecting individuals based on available neuroticism sum-score and active consent for participation, 372,903 individuals remained for the analyses.

To correct for population-stratification, 30 principal components were calculated on the subset of QC-ed unrelated European subjects based on 145,432 independent (*r*^*2*^<0.1) SNPs with MAF>0.01 and INFO=1 using FlashPC2^35^.Subsequently, imputed variants were converted to hard call using a certainty threshold of 0.9. Multi-allelic SNPs, indels, and SNPs without unique rs id were excluded, as well as SNPs with a low imputation score (INFO score <0.9), low minor allele frequency (MAF<0.0001) and high missingness (>0.05). This resulted in a total of 10,847,151 SNPs used for downstream analysis.

#### UK Biobank – Depression

Similar genotyping/imputation/filtering procedures as described above for the UKB neuroticism GWAS were followed for the UKB depression GWAS, resulting in N=362,696.

***Other samples***: Summary statistics were used for 23andMe and PGC. Genotyping and imputation of these samples are described in detail elsewhere (23andMe depression^28^; PGC depression^27^).

### Genome-wide association analyses

#### UK Biobank – Neuroticism

Genome-wide association analyses were performed in PLINK^36,37^, using a linear regression model of additive allelic effects with age, sex, townsend deprivation index, genotype array, and 10 genetic European-based principal components as covariates.

#### UK Biobank – Depression, depressed affect, worry

The settings, covariates, and exclusion criteria for the UKB depression, UKB *depressed affect*, and UKB *worry* GWAS were the same as described above for UKB neuroticism GWAS, with 10,847,151 SNPs remaining after all exclusion steps.

***Other samples***: Summary statistics were used for 23andMe, GPC and PGC. Details on the genome-wide association analyses of these samples can be found elsewhere (23andMe 10 28 9 27 neuroticism ; 23andMe depression ; GPC neuroticism ; PGC depression).

### Meta-analysis

#### Neuroticism

Meta-analysis of the neuroticism GWAS in UKB, 23andMe, and GPC was carried out in METAL^16^. The meta-analysis was performed on the *P*-value of each SNP using a sample size-weighted fixed-effects analysis. Bonferroni correction was applied to correct for multiple testing. The genetic signal correlated strongly between the three samples (*rg* range: 0.83 – 1.07; **Supplementary Table 1**), supporting the decision to meta-analyze.

#### Depression

Meta-analysis of the depression GWAS in UKB, 23andMe and PGC was carried out in METAL^16^. As the UKB GWAS concerned a continuous operationalization of the depression phenotype, while 23andMe and PGC used case-control phenotypes, the odds ratio from the 23andMe and PGC summary statistics were converted to log odds, reflecting the direction of the effect. The meta-analysis was then performed on the *P*-value of each SNP using a sample size-weighted fixed-effects analysis. Bonferroni correction was applied to correct for multiple testing. Genetic correlations between the three samples were moderate to strong (*rg* range: 0.61 – 0.80; **Supplementary Table 22**).

##### Functional Annotation

Functional annotation was performed using FUMA^17^ (http://fuma.ctglab.nl/), an online platform for functional mapping of genetic variants. We first defined independent significant SNPs which have a genome-wide significant *P*-value (5´10^-8^) and are independent at *r^2^*<0.6. Lead SNPs were defined by retaining those independent significant SNPs that were independent from each other at *r*^*2*^<0.1 (based on LD information from UK Biobank genotypes; see **Supplementary Information** for a more detailed explanation). Subsequently, risk loci were defined by merging lead SNPs that physically overlapped or whose LD blocks were closer than 250kb apart. As a result, when analyzing multiple phenotypes, as in the current study, the same locus may be discovered for different phenotypes, whilst different lead SNPs are identified.

We selected all SNPs with *r^2^*>0.6 with one of the independent significant SNPs, a *P*-value lower than 0.05 and minor allele frequency (MAF) higher than 0.0001 for annotations. Functional consequences for all independent significant SNPs and SNPs in LD with them were obtained by performing ANNOVAR gene-based annotation using Ensembl genes. In addition, CADD scores (indicating the deletriousness of SNP, with scores >12.37 seen as likely deleterious)^38^ and RegulomeDB scores (with lower scores indicating a higher probability of having a regulatory function) were annotated to SNPs by matching chromosome, position, reference and alternative alleles.

##### Gene-mapping

SNPs in genomic risk loci that were GWS or were in LD (>0.6) with one of the independent GWS SNPs were mapped to genes in FUMA^19^ using three strategies:

1. Positional mapping maps SNPs to genes based on the physical distances (i.e., within 10kb window) from known protein coding genes in the human reference assembly (GRCh37/hg19).
2. eQTL mapping maps SNPs to genes with which they show a significant eQTL association (i.e. the expression of that gene is associated with allelic variation at the SNP). eQTL mapping uses information from 3 data repositories (GTEx, Blood eQTL browser BIOS QTL browser, and is based on cis-eQTLs which can map SNPs to genes up to 1Mb apart. A false discovery rate (FDR) of 0.05 was applied to define significant eQTL associations.
3. Chromatin interaction mapping was performed to map SNPs to genes based on a significant chromatin interaction between a genomic region in a risk locus and promoter regions of genes (250bp up and 500bp downstream of transcription start site (TSS)). Chromatin interaction mapping can involve long-range interactions as it does not have a distance boundary as in eQTL mapping. FUMA currently contains Hi-C data of 14 tissue types from the study of^39^. Since chromatin interactions are often defined in a certain resolution, such as 40kb, an interacting region may span multiple genes. All SNPs within these regions would be mapped by this method to genes in the corresponding interaction region. To further prioritize candidate genes from chromatin interaction mapping, we integrated predicted enhancers and promoters in 111 tissue/cell types from the Roadmap Epigenomics Project^40^; chromatin interactions are selected in which one region involved in the interaction overlaps with predicted enhancers and the other region overlaps with predicted promoters in 250bp up- and 500bp downstream of TSS site of a gene. We used a FDR of 1×10^-5^ to define significant interactions.

### Gene-based analysis

A genome-wide gene association analysis (GWGAS) can identify genes in which multiple SNPs show moderate association to the phenotype of interest without reaching the stringent genome-wide significance level. At the same time, as a GWGAS takes all SNPs within a gene into account, a gene harbouring a genome-wide significant SNP may not be implicated by a GWGAS analyses when multiple other SNPs within that gene show only very weak association signal. The *P-*values from the SNP-based GWAS meta-analyses for neuroticism and depression, and the GWAS for *depressed affect* and *worry*, were used as input for the genomewide gene association analysis (GWGAS) in MAGMA (http://ctg.cncr.nl/software/magma)^23^, and all 19,427 protein-coding genes from the NCBI 37.3 gene definitions were used. We annotated all SNPs in our GWA (meta-) analyses to these genes, resulting in 18,187, 18,187, 18,182, and 18,182 genes that were represented by at least one SNP in the neuroticism metaanalysis, the depression meta-analysis, the *depressed affect* GWAS, and the *worry* GWAS, respectively. We included a window around each gene of 2 kb before the transcription start site and 1 kb after the transcription stop site. Gene association tests were performed taking into account the LD between SNPs, and a stringent Bonferroni correction was applied to correct for multiple testing (0.05/number of genes tested: *P*<2.75×10^-6^).

### Gene-set analysis

We used MAGMA^23^ to test for association of predefined gene-sets with neuroticism, depression, *depressed affect*, and *worry*. A total of 7,246 gene-sets were derived from several resources, including BioCarta, KEGG, Reactome^41^ and GO. All gene-sets were obtained from the MsigDB version 5.2 (http://software.broadinstitute.org/gsea/msigdb/collections.jsp). In addition, we performed gene-set analysis on 53 tissue expression profiles obtained from the GTEx portal (https://www.gtexportal.org/home/), and 24 cell-type specific expression profiles.

Definition and calculation of gene-sets for cell-type specific expression is described in detail elsewhere^26,42^. Briefly, brain cell-type expression data was drawn from scRNAseq data from mouse brain^26^. For each gene, the value for each cell-type was calculated by dividing the mean Unique Molecular Identifier (UMI) counts for the given cell type by the summed mean UMI counts across all cell types^26^. Associations between gene-wise *P*-values from the meta-analysis and cell-type specific gene expression were calculated using MAGMA^23^, by grouping genes into 40 equal bins by specificity of expression, and regressing bin-membership on gene-wise association with neuroticism in the meta-analysis. Results were considered significant if the association *P*-values were smaller than the relevant Bonferroni threshold.

For all gene-sets we computed competitive *P-*values, which result from the test whether the combined effect of genes in a gene-set is significantly larger than the combined effect of a same number of randomly selected genes (in contrast, self-contained *P*-values result from testing against the null hypothesis of no effect). We only report competitive *P*-values, which are more conservative compared to self-contained *P*-values. Competitive *P*-values were Bonferroni corrected (α=0.05/7,323=6.83´10^-6^).

Conditional gene-set analyses were performed with MAGMA as a secondary analysis to test whether each observed enriched cell-type was independent of all others. Full details of the method implemented are provided in ^26^.

### Genetic correlations

Genetic correlations (*r_g_*) were computed using LDscore regression^17,18^ (https://github.com/bulik/ldsc). The significance of the genetic correlations of neuroticism, depression, *depressed affect* and *worry* with 35 behavioral, social and (mental) health phenotypes for which summary statistics were available was determined while correcting for multiple testing through a stringent Bonferroni corrected threshold of *P*<0.05/(4×35) (3.6×10^-4^).

### Partitioned heritability

To investigate the relative contribution to the overall heritability of SNPs annotated to 22 specific genomic categories, we partioned SNP heritability by binary annotations using stratified LD score regression^20,43^. Information about binary SNP annotations were obtained from the LD score website (https://github.com/bulik/ldsc). Enrichment results reflect the X-fold increase in *h*^2^ proportional to the number of SNPs (e.g., enrichment=13.79 for SNPs in conserved regions implies that a 13,79-fold increase in *h*^2^ is carried by SNPs in these region, corrected for the proportion of SNPs in these regions compared to all tested SNPs).

### Polygenic risk scoring

To test the predictive accuracy (Δ*R*^2^) of the our meta-analytic results, we calculated a polygenic risk score (PGS) based on the SNP effect sizes of the current analysis. As independent samples we used holdout samples; we removed 3,000 individuals from the discovery sample (UKB only, as we only had access to raw data from this sample) and reran the genome-wide analyses. We repeated this three times, to create 3 randomly drawn, independent hold-out samples. Next, we calculated a PGS on the individuals in each of the 3 holdout samples. PGS were calculated using LDpred^22^ and PRSice^21^ (clumping followed by *P*-value thresholding).

For LDpred, PGS were calculated based on diferent LDpred priors (*P_LDpred_* = 0.01, 0.05, 0.1, 0.5, 1 and infinitesimal). The explained variance (*R*^2^) was derived from the linear model, using the neuroticism summary score as the outcome, while correcting for age, gender, array, batch and genetic principal components.

### Data availablity

Our policy is to make genome-wide summary statistics (sumstats) publically available. Sumstats from our neuroticism meta-analysis, our depression meta-analysis, and the GWA analyses for *depressed affect* and *worry* are available for download at https://ctg.cncr.nl/.

Note that our freely available meta-analytic sumstats concern results excluding the 23andMe sample. This is a non-negotiable clause in the 23andMe data transfer agreement, intended to protect the privacy of the 23andMe research participants. To fully recreate our meta-analytic results for neuroticism: (a) obtain Lo et al. (2016) sumstats from 23andMe (see below); (b) conduct a meta-analysis of our sumstats with the Lo et al. sumstats. To fully recreate our metaanalytic results for depression: (a) obtain Hyde et al. (2016) sumstats from 23andMe (see below); (b) conduct a meta-analysis of our sumstats with the Hyde et al. sumstats.

23andMe participant data are shared according to community standards that have been developed to protect against breaches of privacy. Currently, these standards allow for the sharing of summary statistics for at most 10,000 SNPs. The full set of summary statistics can be made available to qualified investigators who enter into an agreement with 23andMe that protects participant confidentiality. Interested investigators should contact David Hinds for more information.

